# PDZ Domains from the Junctional Proteins Afadin and ZO-1 Act as Mechanosensors

**DOI:** 10.1101/2023.09.24.559210

**Authors:** Vipul T. Vachharajani, Matthew P. DeJong, Soumya Dutta, Jonathan Chapman, Eashani Ghosh, Abhishek Singharoy, Alexander R. Dunn

## Abstract

Intercellular adhesion complexes must withstand mechanical forces to maintain tissue cohesion while also retaining the capacity for dynamic remodeling during tissue morphogenesis and repair. Many cell-cell adhesion complexes contain at least one PSD95/Dlg/ZO-1 (PDZ) domain situated between the adhesion molecule and the actin cytoskeleton. However, PDZ-mediated interactions are characteristically nonspecific, weak, and transient, with multiple binding partners per PDZ domain, micromolar dissociation constants, and bond lifetimes of seconds or less. Here, we demonstrate that the bonds between the PDZ domain of the cytoskeletal adaptor protein afadin and the intracellular domains of the adhesion molecules nectin-1 and JAM-A form molecular catch bonds that reinforce in response to mechanical load. In contrast, the bond between the PDZ3-SH3-GUK (PSG) domain of the cytoskeletal adaptor ZO-1 and the JAM-A intracellular domain becomes dramatically weaker in response to ∼2 pN of load, the amount generated by single molecules of the cytoskeletal motor protein myosin II. Thus, physiologically relevant forces can exert dramatic and opposite effects on the stability of two of the major linkages between cell-cell adhesion proteins and the F-actin cytoskeleton. Our data demonstrate that that PDZ domains can serve as force-responsive mechanical anchors at cell-cell adhesion complexes. More broadly, our findings suggest that mechanical force may serve as a previously unsuspected regulator of the hundreds of PDZ-ligand interactions present in animal cells.

## Introduction

The construction of epithelial tissues requires the coordination of multiple classes of transmembrane adhesion molecules that bind across the cell-cell junction to connect neighboring cells (**Fig. 1A**)^1,2^. These linkages between cells must maintain robust adhesion in the face of mechanical forces due to both external stresses and cellular events such as division and death, while also maintaining the dynamic properties that make tissue remodeling and repair possible^3–5^. How these seemingly contradictory properties are achieved at the molecular level is in general poorly understood. Previous studies have examined how the components of cadherin-mediated adherens junctions respond to mechanical load^6–13^. In contrast, less is known about how the other adhesion complexes present at cell-cell junctions may respond to force^14^.

**Figure 1.**
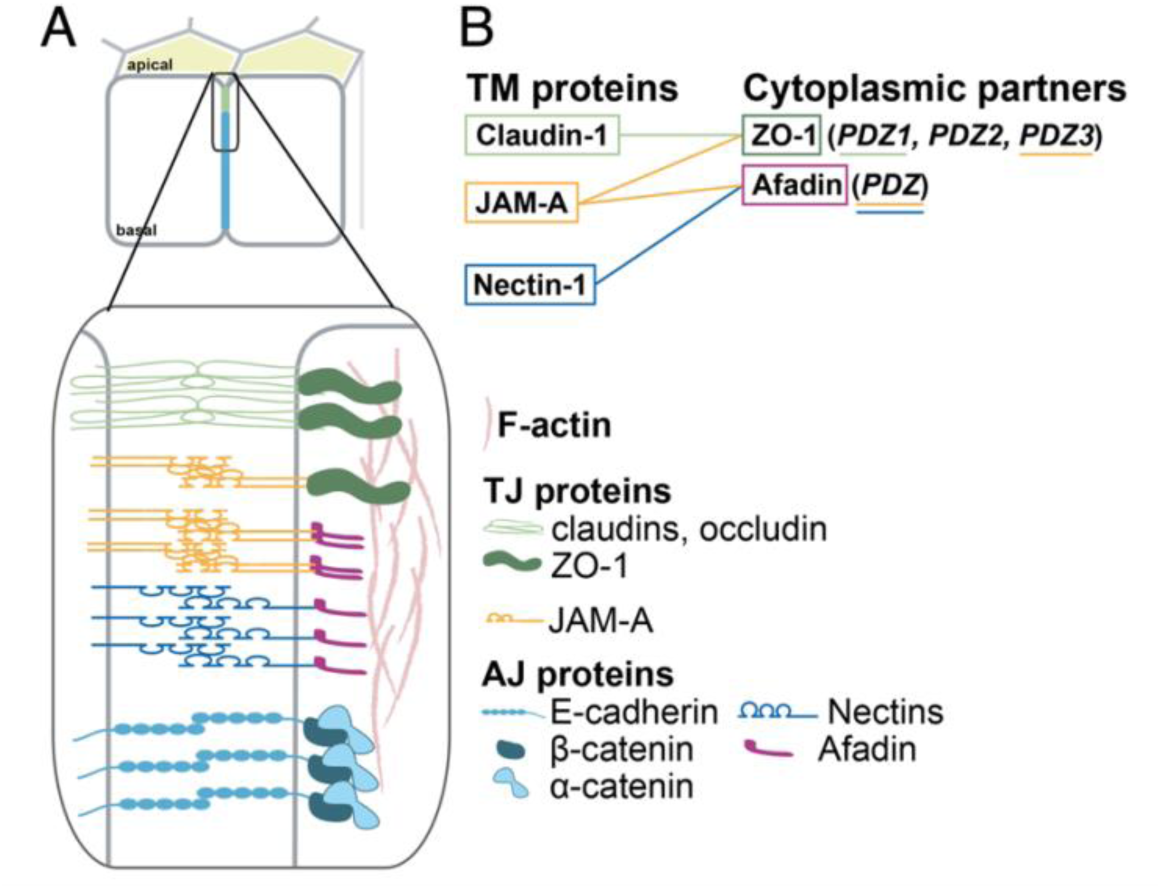
Apicobasal organization of cell-cell junctions. (A) Schematic showing the diversity of cell-cell adhesion molecules and intracellular scaffold proteins at adherens and tight junctions (AJs, and TJs, respectively). Modified with permission from Vasquez, de la Serna, and Dunn (2021). (B) Simplified network of PDZ-mediated interactions between transmembrane (TM) proteins and their cytoplasmic binding partners.

Unlike the cadherin-catenin complex, most other cell-cell adhesion complexes contain a cytoskeletal linker protein containing at least one PDZ domain^1^. In general, PDZ domains bind to the C-terminal ∼5 residues of multiple binding partners, resulting in a complex network of PDZ-peptide interactions (**Fig. 1B**)^15–18^. Afadin and zonula occludens-1 and -2 (ZO-1/2) are PDZ containing scaffolding proteins that help to organize adherens and tight junctions at cell-cell contacts, respectively^1,19–22^. Both proteins bind transmembrane cellular adhesion proteins via their PDZ domains and link directly to filamentous, (F)-actin, providing a potential conduit for force transmission between cell adhesion complexes and the actin cytoskeleton. ZO-1, afadin, and their binding partners are known to be mechanoresponsive, such that mechanical tension exerted at cell-cell junctions leads to altered intracellular signal transduction and transcriptional regulation^14,23–28^. However, to our knowledge how and whether PDZ-mediated linkages by afadin, ZO-1, or any other protein may respond to mechanical load remains unexamined.

It is not clear how the intricate organization of cell-cell junctions can arise from an apparently promiscuous network of overlapping interactions. Afadin possesses one PDZ domain that binds to either the JAM family of tight junction proteins or to the nectin family of adherens junction proteins (**Fig. 1**)^29,30^. ZO proteins have three PDZ domains. Of these, PDZ1 binds to claudins, while PDZ3 binds to JAMs^31–33^. Importantly, PDZ3 exists as part of structural module containing PDZ, Src homology 3 (SH3), and guanylate kinase (GUK) domains, termed the PSG module^32,34^. The ZO-1 PSG forms a dense web of interactions with the cytoskeletal adaptor proteins afadin, αE-catenin, vinculin, and shroom2, signal transduction proteins such as ZONAB and Gα_12_, and the tight junction protein occludin^23,34–42^. In addition to its central role in the construction of tight junctions, ZO-1 plays an integral role in the formation of new cell-cell contacts and is recruited to nascent adherens junctions through its interactions with afadin and αE-catenin^43–48^. The factors that regulate ZO-1 localization of tight vs. adherens junctions, and more broadly its interactions with a diverse array of binding partners, remain poorly understood.

## Results

### Nectin-1 and JAM-A form short-lived, µM-affinity bonds with the PDZ domain of afadin in solution

Unusually high affinities of the afadin PDZ domain for nectin and JAM-A could potentially explain the function of this linkage as a load-bearing linkage to the actin cytoskeleton. To directly measure affinities of afadin PDZ-binding ligands, we first measured the solution dissociation constants and binding kinetics of the intracellular domains (ICDs) of nectin-1 and JAM-A with the afadin PDZ domain (**Fig. 2A, B**). The affinity of the nectin-1 ICD with the afadin PDZ was approximately 3 µM as measured by both bio-layer interferometry (**Fig. 2C**) and fluorescence anisotropy (**Supp. Fig. 1A**). The affinity of the JAM-A ICD and the afadin PDZ was likewise approximately 5 µM (**Fig. 2C, Supp. Fig. 1B**). These values that are consistent with previous measurements of PDZ-ligand dissociation constants.

**Figure 2.**
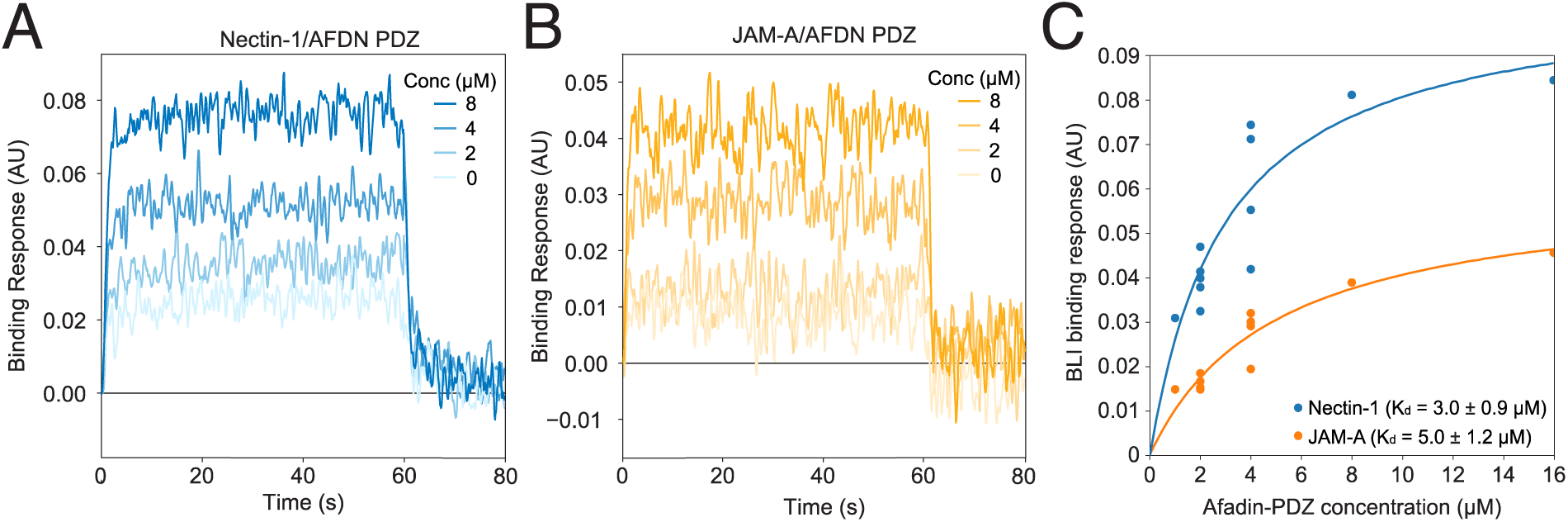
JAM-A and nectin-1 ICDs have μM affinities for the afadin PDZ and dissociate within seconds. Representative BLI responses of (A) the nectin-1 ICD binding to the Afadin PDZ domain and (B) the JAM-A ICD binding to the afadin PDZ domain. (C) Equilibrium binding data for nectin-1 ICD/afadin PDZ and JAM- A ICD/Afadin PDZ indicate K_D_ values of 3 and 5 µM, respectively.

We next tested the hypothesis that the bonds of the afadin PDZ to JAM-A, nectin-1, or both might be unusually long-lived. We measured binding kinetics using bio-layer interferometry and found that both ligands formed bonds with lifetimes on the single-second timescale, with bond lifetimes of 1.2 and 0.65 s for the nectin-1 and JAM-A ICDs, respectively (**Fig. 2A, B, Supp. Fig. 2C**). These relatively modest dissociation constants and bond lifetimes are striking given the hypothesized role of nectins in anchoring F-actin at cell-cell junctions^49–51^. In addition, the similarity in binding lifetimes suggests that it is unlikely that this provides a means by which afadin may distinguish between these two binding partners.

### Afadin-PDZ forms catch bonds with C-terminal ligands under force

We next used a single-molecule magnetic tweezers assay to measure the lifetimes of bonds between the afadin PDZ and nectin-1 and JAM-A ICDs under physiologically relevant, piconewton forces. To do so, we designed a tethered-ligand construct in which the PDZ domain of afadin is linked to the C-terminal domain of either nectin-1 or JAM-A by a flexible glycine/serine linker, with handles for stable attachment on either side (**Fig. 3A**). These constructs were covalently attached on one end to a glass coverslip functionalized with a SpyCatcher-conjugated elastin-like polypeptide (ELP) and tethered to a 2.7 µm paramagnetic bead on the other end via a biotin-streptavidin interaction.

**Figure 3.**
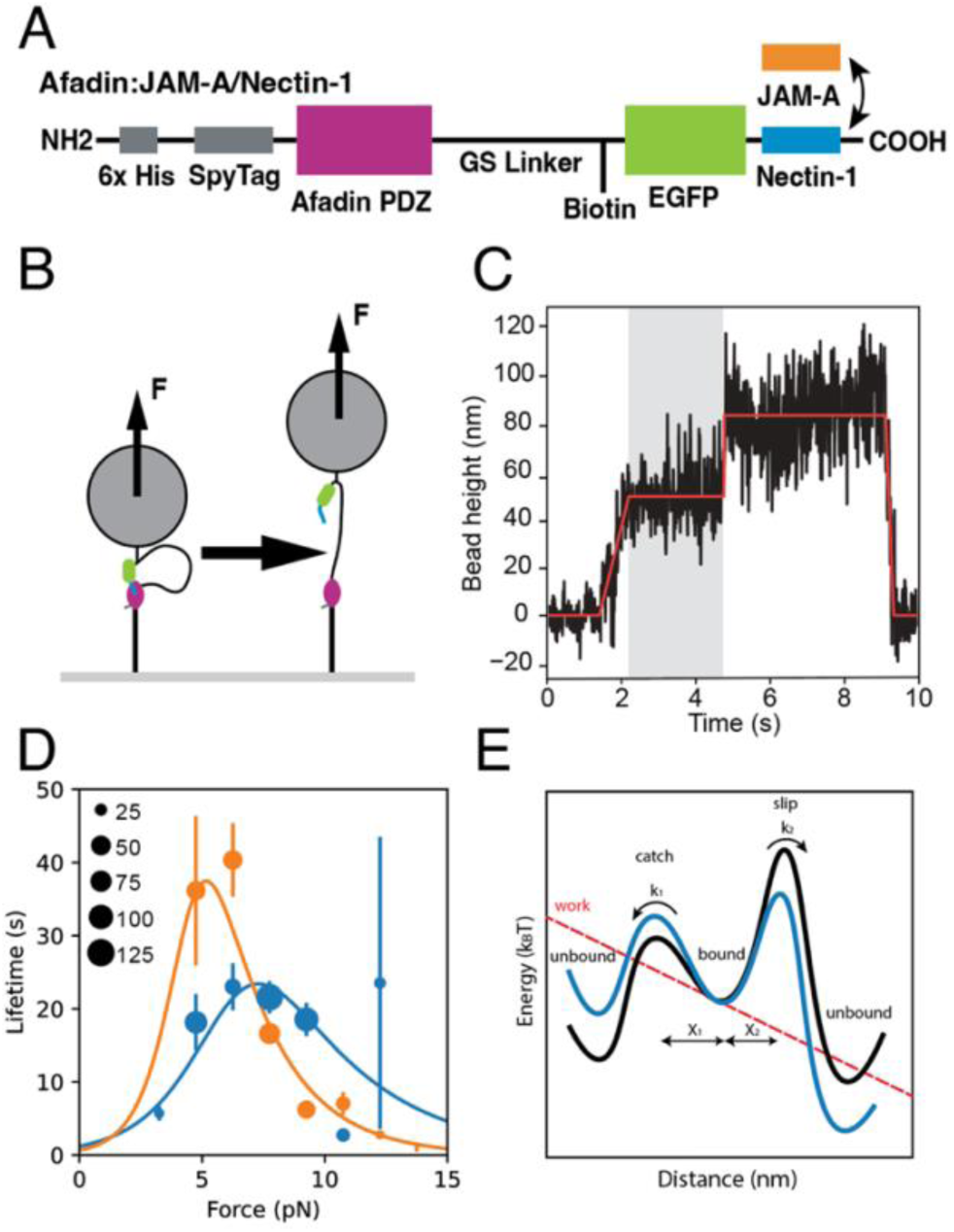
A single-molecule magnetic tweezers assay to measure afadin PDZ unbinding kinetics under tension. (A) Domain schematics for the magnetic tweezers afadin PDZ/nectin-1 or JAM-A fusion constructs. (B) Schematic of the force-jump magnetic tweezer experiment performed on tethered protein constructs. (C) Sample magnetic tweezers trace, plotting bead height vs. time. Actual data are shown in *black*, and a two-step fit is shown in *red*. The shaded region, during which force is applied but the linker has not extended, corresponds to the measured bond lifetime. (D) Average bond lifetime vs. force for the nectin-1 ICD (*blue*, *N* = 801) and JAM-A ICD (*orange*, *N* = 567) constructs. Marker size corresponds to the number of observations. The open circles represent measured bond lifetimes at zero force from BLI. Filled circles correspond to the means of 1.5 pN force bins. Catch bond fits, constrained by the zero-force lifetime, are shown in solid lines for nectin-1/afadin and JAM-A/afadin. Error bars denote the standard deviation of bond lifetimes for each bin. (E) Schematic for the catch bond model. The PDZ-peptide complex dissociates via catch (*k_1_*) and slip (*k_2_*) pathways. Mechanical load results in a mechanical work term (force × distance; *red* dotted line) that tilts the energetic landscape (*blue*), slowing *k_1_*, and accelerating *k_2_*. Distances to the transition states are indicated by *x_1_* and *x_2_*.

To test the hypothesis that force modulates afadin-PDZ bond kinetics, we performed force-jump experiments. Briefly, bead height was monitored as the force on the bead was increased from <1 pN to a fixed force between 3 and 15 pN (**Fig. 3B**). The time between the end of the initial force ramp and the steplike change in bead height corresponding to linker extension was taken as the bond lifetime (**Fig. 3C**). To confirm that the step we measured corresponded to the appropriate linker extension, we repeated the experiment using a construct containing a longer linker and observed a corresponding increase in step height (**Supp. Fig. 3**).

As force increased from 0 pN, we observed an increase in bond lifetime up to a peak value around 5-7 pN for both nectin-1/afadin and JAM-A/afadin (**Fig. 3D**), after which point the bond lifetime decreased with increasing force. This behavior is consistent with the presence of a molecular catch bond, in which binding lifetimes increase over a given force range. Catch bonds can be described by several kinetic models^52^. Of these, a two-pathway model described the data well (**Fig. 3D; Supp. Fig. 4; Table 1**)^53,54^. In this model, the peptide-PDZ complex can dissociate by one of two pathways. Force accelerates dissociation by one pathway but hinders dissociation by the other (**Fig. 3E**). These results suggest that afadin can form catch bonds with its C-terminal ligands, and thus stably associate with ligands under tension, despite its modest affinity for these ligands in solution.

**Table 1.** Maximum likelihood estimate parameters for catch-bond models. *k*_i_ ^0^ is the zero-force transition rate. *x_i_* is the distance to the corresponding transition state. Parameter uncertainties are reported as the 95% confidence interval (CI).

| **Afadin/Nectin-1** |  |  |  |  |
| --- | --- | --- | --- | --- |
| Transition | $k_i^0$ (1/s) | CI (1/s) | $x_i$ (nm) | CI (nm) |
| $k_1$ | 0.829 | [0.828, 0.830] | 2.30 | [2.28, 2.35] |
| $k_2$ | 0.0041 | [0.0035, 0.0053] | 1.09 | [0.99, 1.15] |
| **Afadin/JAM-A** |  |  |  |  |
| Transition | $k_i^0$ (1/s) | CI (1/s) | $x_i$ (nm) | CI (nm) |
| $k_1$ | 1.536 | [1.536, 1.536] | 4.20 | [4.16, 4.28] |
| $k_2$ | 0.0022 | [0.0021, 0.0025] | 1.71 | [1.65, 1.74] |
| **ZO-1/JAM-A** |  |  |  |  |
| Transition | $k_i^0$ (1/s) | CI (1/s) | $x_i$ (nm) | CI (nm) |
| $k_1$ | 0.025 | [0.017, 0.039] | 10.3 | [9.5, 11.0] |

### Molecular dynamics simulations offer insight into the possible mechanistic origins of the afadin/nectin-1 catch bond

A combination of equilibrium and biased molecular dynamics (MD) simulations were performed to identify the key residues that collectively contribute to catch bond formation^55^. Although the pulling speed in steered MD is orders of magnitude faster than in magnetic tweezer assays, MD can still offer insights into the non-equilibrium molecular interactions that lead to the catch bond (**Fig. 4**)^56,57^. Here, we performed 14 100 ns steered MD simulations^58,59^ in which the afadin PDZ and nectin-1 C-terminal peptide were subject to a pulling rate of 0.5 Å ns^-1^. These data were then combined to generate an ensemble that samples the equilibrium configurations along this pulling trajectory (see Methods).

**Figure 4.**
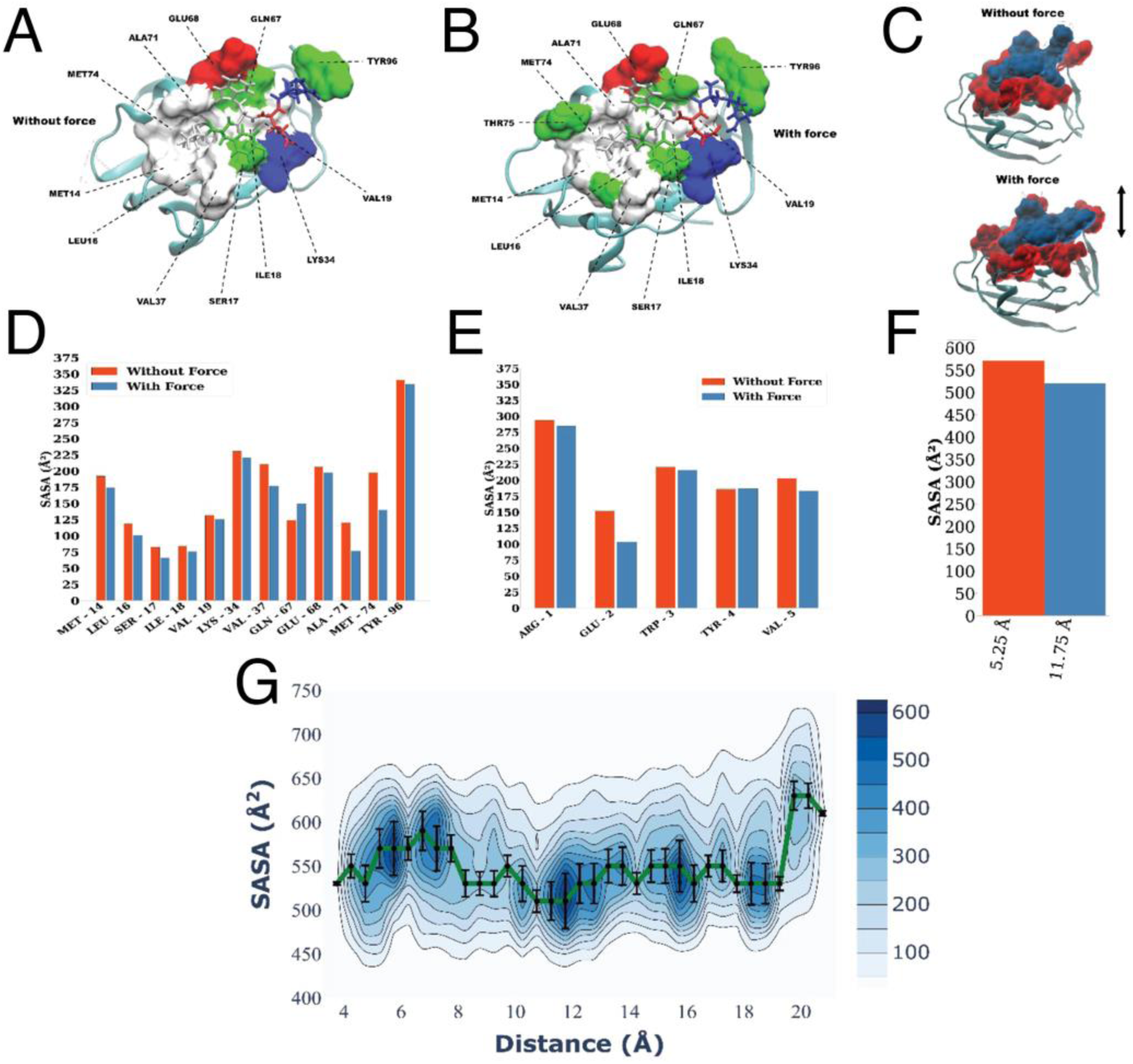
Analysis of afadin/nectin-1 interface from SMD simulations. (A) Surface representation of the residues of the afadin PDZ domain that contact the nectin-1 ICD (shown as sticks) in the absence of imposed force. Coordinates were extracted from structures that populate the most probable region of the SASA distribution from the equilibration run. Hydrophobic residues are shown in *white*, polar residues in *green*, and negative and positive residues are in *red* and *blue* respectively. (B) Structure generated as in (A) illustrating a conformation under simulated force. (C) Total surface representation of the interacting residues of the afadin PDZ (*red*) and nectin ICD (*blue*). The direction of pulling during SMD is shown with a double-headed arrow. (D) Residue-wise SASA comparison for the afadin PDZ with force and without force. (E) Residue-wise SASA comparison for the residues in the nectin ICD, numbered starting with the fifth residue from the C-terminus. In general, lower SASA corresponds to decreased free energy. (F) Total contact area SASA with and without force. (G) A contour plot depicting the SASA around the nectin-1 IDC versus the distance between the C-terminal nitrogen atom of afadin and the N-terminal nitrogen atom of the five-residue nectin peptide, with data density shown in blue color scale, with a maximum SASA probability pathway overlaid in *green*. The vertical bars (*black*) depict the spread of high probability points within 5% around the maximum probability line.

**Figure 5.**
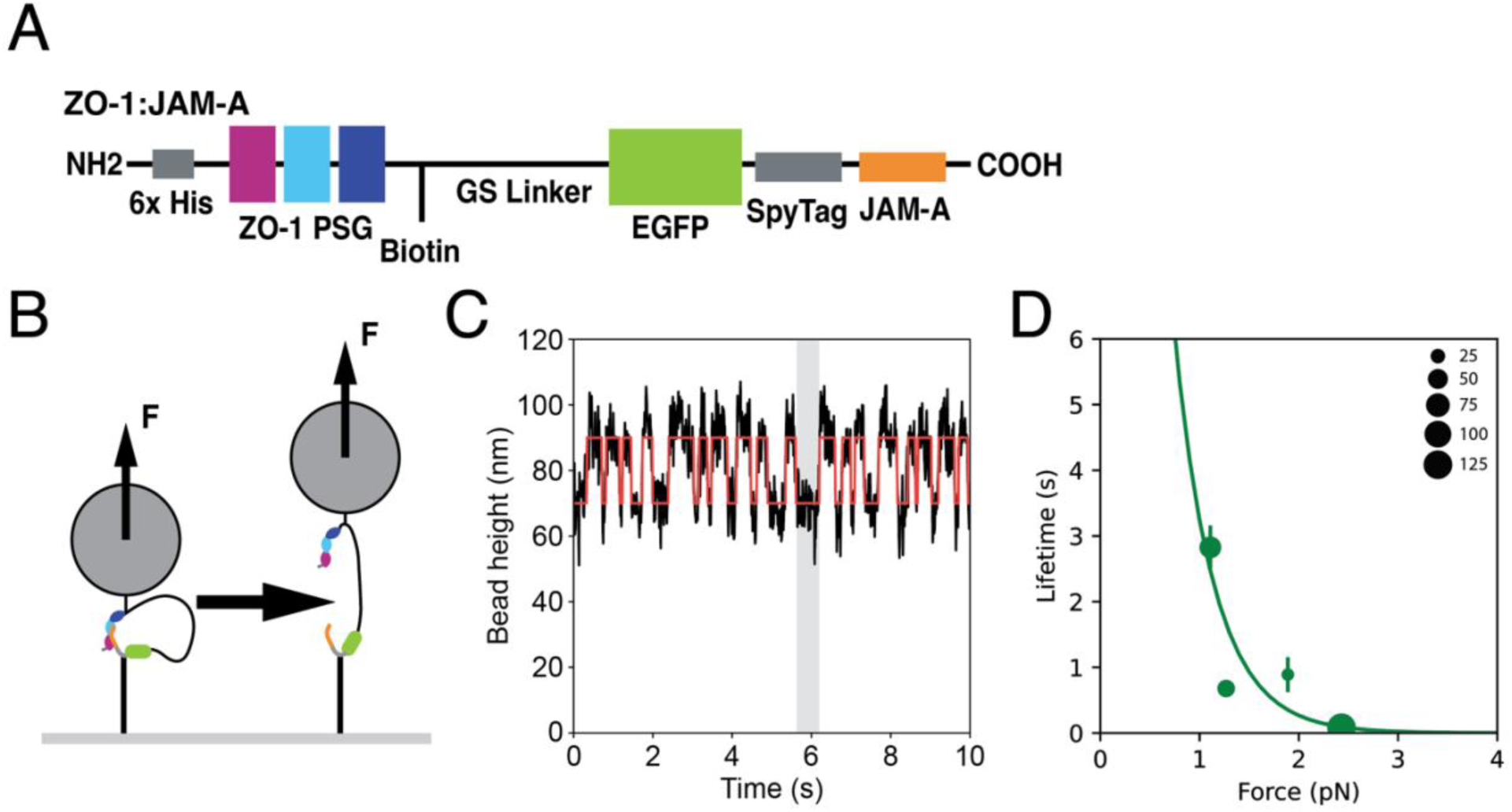
A single-molecule magnetic tweezers assay to measure ZO-1 PSG unbinding kinetics under tension. (A) Domain schematic of the magnetic tweezers ZO-1 PSG/JAM-A fusion construct. The ZO-1 PSG includes a PDZ domain (*pink*), an SH3 domain (*light blue*), and a GUK domain (*dark blue*). (B) Schematic of the force-clamp magnetic tweezer experiment performed on the tethered protein construct. (C) Sample magnetic tweezer trace, plotting bead height over time. Actual data are shown in *black*, and two-step fit in *red*. (D) Average bond lifetime vs force for JAM-A/ZO-1 (*N* = 245). Marker size corresponds to the number of observations. A one-state slip bond fit is shown. Error bars denote standard deviation of bond lifetime for each bin.

We then used simulation data to create a two-dimensional (2D) histogram of the Solvent Accessible Surface Area (SASA) of the interface between the afadin PDZ domain and nectin-1 ICD as a function of the distance between the C-terminal nitrogen atom of the afadin PDZ and the and C-terminal nitrogen atom of nectin-1 ICD (**Fig. 4**). These data show that across multiple, uncorrelated trajectories, the most probable SASA decreases from 570.5 Å^2^ to 520 Å^2^ as the distance between the C-terminal nitrogen atom of afadin and the C-terminal nitrogen atom of the five-residue nectin peptide increases from 5.25 to 11.75 Å. Put differently, the contact area between the afadin PDZ domain and nectin-1 ICD, increases when force is applied, indicating that the complex slides into a more compact conformation due to the application of force. The most probable pathway along the 2D profile reveals that the interface of the afadin PDZ and the nectin-1 ICD packs more efficiently around the C-terminal valine of the nectin-1 ICD. Additionally, force leads to a stabilization of the salt-bridge interaction between Lys34 of the afadin PDZ and Glu2 of the nectin ICD as indicated by decreased SASA for both residues (**Fig. 4D, E**).

### Force accelerates the dissociation of the complex between the JAM-A ICD and the ZO-1 PSG domain

To assess how force selects for certain PDZ-peptide binding interactions, we measured JAMA-ICD dissociation from the ZO-1 PSG domain under force. We designed a tethered-ligand construct in which the PSG fragment of ZO-1 is linked to the C-terminal domain of JAM-A using the same linkers and handles as in the JAM-A/afadin PDZ measurements (**Fig. 4A**). We found that binding was reversible at low pN forces, so we performed experiments in which force is held at a constant value to quantify the effect of load on bond lifetimes (**Fig. 4B, C**). Strikingly, we observed a strong monotonic decrease in bond lifetime to <0.1 s at 2.3 pN (**Fig. 4D**). These data are well-described by a slip bond, in which mechanical force accelerates dissociation from a single bound state (Table 1). The measured *x*_10_, the distance to the transition state for the dissociation step, was 10.3 nm. This large distance implies a correspondingly large conformational rearrangement in the PSG during dissociation (see Discussion).

## Discussion

Our results demonstrate that the afadin PDZ and the ICDs of nectin-1 and JAM-A form bonds that dramatically strengthen under tension despite the relatively short lifetimes of these interactions in solution. In contrast, the ZO-1 PSG domain forms a remarkably force-sensitive slip bond with the JAM-A ICD, whose binding lifetime decreases >40-fold in response to the force generated by one myosin II motor domain. Thus, physiologically relevant forces can exert dramatic and opposite effects on the stability of two of the major linkages between cell-cell adhesion proteins and the F-actin cytoskeleton. Though we examined only two PDZ domains in detail, the abundance of PDZ domains present at cell-cell junctions suggests that PDZ-domain mechanosensitivity may represent a new class of mechanical behaviors that govern the organization and dynamics of cell adhesion ligands.

The catch bond between the afadin PDZ domain and its cognate peptide ligands is unexpected. Several previously proposed models can potentially explain this observation. (1) Force may cause a switch in the conformation of the complexes from a “weakly bound” to a “strongly bound” state, as has been proposed for the interactions of talin, vinculin, and αE-catenin with F-actin, and the interaction of von Willebrand factor with glycoprotein Ib.^7,60–63^ (2) Force may cause a *<u>continuous</u>* change in conformation that strengthens the interaction with increasing force due to force-driven changes in the binding interface. This model has likewise been proposed to describe catch bonding by P and L selectins.^64^ (3) Alternatively, partial unfolding in the transition state has been proposed to impose an entropic penalty that, in some cases, can result in catch bond behavior.^65^ (4) Finally, force can potentially bias dissociation between two discrete pathways (*i.e.* Fig. 3E). Model (1) is not consistent with the monoexponential bond lifetime distributions observed for the JAM-A and nectin-1 ICDs bound to the afadin PDZ (Fig. S5, S6) (ref). Models (2) - (4) predict monoexponential lifetime distributions at a given force, and are thus in principle consistent with our data. Of these, a two-pathway catch bond mechanism, i.e. (3), yields the best fit as judged by the Akaike information criterion (AIC, *not shown*).^66^ Importantly, the physical mechanisms underlying models (2) - (4) are not necessarily mutually exclusive. Simulations additionally suggest a deformation of PDZ domain in response to load that can potentially strengthen binding (**Fig. 4**). The molecular mechanism underlying this PDZ-peptide catch bond thus remains to be determined and constitutes an interest avenue for future investigations.

The large value of *x_1_* for the dissociation of the ZO-1 PSG and JAM-A ICD implies that a large structural rearrangement is coupled to dissociation. Previous experiments demonstrate that both the PDZ and SH3 domains are required for detectable binding between the ZO-1 PSG and JAM-A ICD, and the structure of the complex reveals contacts between the JAM-A ICD and the SH3 domain^32^. A 39-residue disordered loop connects the SH3 and GUK domains. Extension of this loop and rotation of the GUK domain would yield an increase in overall length of ∼15 nm, in good agreement with the 10 nm extension implied by the value of *x*_1_. We speculate that undocking of the PDZ, SH3, and GUK domains precedes dissociation of the JAM-A ICD under load. If so, such a mechanism would explain the remarkable force sensitivity of the dissociation step. These data raise the intriguing possibility that mechanical force may regulate the interaction of ZO-1 with the numerous structural and signaling proteins that bind to the PSG domain.^23,34–42^

It is likely that nectins play an important role in the initiation of new cell-cell contacts. Nectin *trans*-heterodimers have nM affinity^67,68^, much stronger than the µM affinity of the cadherin extracellular domain^69^. This and other observations have led to the hypothesis that nectin- and afadin-based adhesions may form first and then template cadherin-based adhesions^36,49,70,71^. Our results suggest a potential mechanism by which initial nectin-only contacts may robustly recruit afadin to perform this role. Further experiments are necessary to more directly test the hypothesis that *trans-*interacting nectins and afadin bear tension in cells.

Two nectinopathies have been identified in humans: a nectin-1 dominant negative mutation is associated with the cleft lip/palate-ectodermal dysplasia syndrome (CLPED), and a nectin-4 deletion in the afadin PDZ-binding C-terminus has been associated with an ectodermal dysplasia/cutaneous syndactyly syndrome (EDSS1)^72–76^. Furthermore, in a murine model, an afadin conditional knockout phenocopies a nectin-1/nectin-4 double knockout, resulting in the same phenotype as observed in humans with CLPED^77^. These results clearly identify the nectin-afadin linkage as instrumental to ectodermal morphogenesis, and it is intriguing to speculate that a PDZ catch bond may contribute to this important role.

A growing body of evidence implicates JAM-A in cellular mechanosensing^14,24,25^. In one such example, application of tension to the extracellular domain of JAM-A led to the activation of RhoA downstream of serine phosphorylation of the JAM-A ICD^24^. Other studies indicate that JAM-A indirectly regulates cytoskeletal contractility at cell-cell junctions through p114RhoGEF and PDZ-GEF1, and the G protein Rap2^25,78,79^. Understanding how these, and likely other, PDZ-mediated forms of mechanosensing contribute to junctional mechanotransduction offers a promising avenue for future investigations.

Recent studies support a model in ZO-1 provides a liquid-like, fluid interface between tight junctions and F-actin^80^. Our data suggest that the ZO-1-JAM-A linkage is particularly sensitive to applied force, analogous to shear-thinning liquids whose viscosity decreases in response to applied load, a property that may allow the tight junction to remodel as cell-cell junctions change in length. We note as well that the relative affinities of JAM-A for ZO-1 and afadin are predicted to change in opposite directions as a function of mechanical load. We therefore speculate that mechanical force may play a key role and previously unrecognized role in regulating the complex web of interactions that underlie the intricate organization of epithelial cell-cell junctions.

## Acknowledgements

V.T.V. is supported by the Stanford Medical Scientist Training Program (NIH T32GM007365) and the NIH NIDDK (1F30DK124985). M.P.D is supported by the NSF Graduate Research Fellowship (DGE-1656518). A.R.D. acknowledges the HHMI (Faculty Scholar Award) and the NIH (R35GM130332) for funding that supported this research.

## Methods

### Plasmids, protein expression, and purification

A plasmid encoding the GST-tagged human afadin PDZ domain was a gift from Sachdev Sidhu (Addgene plasmid # 103938 ; http://n2t.net/addgene:103938 ; RRID:Addgene_103938). Plasmids encoding tethered fusion constructs were custom cloned by Epoch Biosciences (Missouri City, TX). Proteins were expressed in *E. coli* BL21(DE3) chemically competent cells (New England Biolabs). Cultures of 500 mL in LB broth were induced at OD 0.6 with 1mM IPTG at 18 °C overnight. The cultures were then spun down at 6000×G for 30 minutes.

For GST-tagged proteins, bacterial pellets were resuspended in PBS pH 7.5 with a cOmplete EDTA-free protease inhibitor cocktail (11873580001, Roche), 70 μg/mL lysozyme (90082, ThermoFisher), and 7 U/mL DNase I (04536282001, Roche). The resuspended cells were rotated end-over-end for 30 min at 4 °C, lysed with a tip sonicator, and spun at 12,000×G for 30 minutes to pellet bacterial cell debris. The supernatant was clarified by filtering through a 0.44 ×G μm PVDF syringe filter and then incubated with 2 mL of Pierce glutathione agarose resin (16100, ThermoFisher) for 2 hr at 4 °C in a 5 mL polypropylene column. The resin was washed with four column volumes of PBSTR buffer (PBS, 1M NaCl, 0.005% Tween-20, 1mM DTT) followed by four column volumes of HRV-3C protease cleavage buffer. The column was stopped, and one column volume of cleavage buffer added along with 6x His-tagged HRV-3C protease (kind gift from Elise Bruguera, Stanford University) to remove the GST tag. The column was capped and rotated overnight at 4 °C and the eluate collected. The eluate was incubated with Ni:NTA coated magnetic beads (MyOne HisPur, ThermoFisher) and separated with a magnet to remove residual 3C protease, and then concentrated using a 3 kDa MWCO spin column (Amicon). Protein was divided into 20 μL aliquots, flash frozen in liquid nitrogen, and stored at −80 °C until use.

For His-tagged proteins, pelleted cells were resuspended lysis buffer (50 mM sodium phosphate, 300 mM NaCl, and 10 mM imidazole, pH 8) with cOmplete EDTA-free protease inhibitor cocktail (11873580001, Roche), 70 μg/mL lysozyme (90082, ThermoFisher), and 7 U/mL DNase I (04536282001, Roche). Cells were lysed and the lysate clarified as above, then incubated with 2 mL HisPur resin at 4 °C for 2 hours. The solution was then packed into a gravity column, washed three times with 5 mL of wash buffer (50 mM sodium phosphate, 300 mM NaCl, and 20 mM imidazole, pH 7.4 with 2 mM β-mercaptoethanol; BME), and protein was eluted by incubating the resin bed with 1 mL of elution buffer (50 mM sodium phosphate, 300 mM sodium chloride, and 250 mM imidazole, pH 7.4 and 2 mM BME) for 5 minutes, four times. The eluate was collected, concentrated, and buffer-exchanged into storage buffer (phosphate-buffered saline, 2 mM BME) using Amicon centrifugal filter units (MilliporeSigma) of the appropriate molecular-weight cutoff, then flash-frozen and stored at −80 °C.

### Zero-force binding assays

All binding experiments were performed in buffer containing 50 mM HEPES, 150 mM NaCl, 1 mg/mL Bovine Serum Albumin, and 0.02% Tween-20 at pH 7.5.

#### Protein expression and purification

The C-terminal 10mer peptides of nectin-1 and JAM-A (sequences in Supplementary Table 1) were purchased lyophilized from Genscript with N-terminal biotinylation and N-terminal FITC conjugation. nectin-1 peptide was redissolved at 1 mg/mL concentration in MilliQ water; JAM-A peptide was redissolved at 10mg/mL in anhydrous DMSO.

#### Bio-layer interferometry

Bio-layer interferometry was conducted using the GatorPrime instrument (Gator Bio, Palo Alto, CA), using streptavidin-coated probes. Probes were loaded with 100 ng/mL biotinylated peptide for 4 min, then equilibrated in buffer for 10 minutes. Binding kinetics were measured at various concentrations of afadin PDZ using an association phase of 300 s and a dissociation phase of 600 s after dipping into fresh wells of buffer. Repeated association and dissociation curves were obtained sequentially from the same probe at 2 mM and 4 mM concentrations. Experiments were repeated on three separate days with separate probes. A blank probe without peptide was used to correct for nonspecific adsorption of PDZ to the probe.

#### Fluorescence anisotropy

Fluorescence anisotropy was measured on a BioTek Synergy 2 plate reader. Briefly, FITC-labeled peptide at a final concentration of 50 ng/mL was mixed with the afadin PDZ domain of varying concentrations in black low-binding 96-well plates (Costar).

### Magnetic tweezers

#### Construction and calibration

A magnetic tweezer was constructed on a Nikon Ti-E microscope equipped with a motorized 3- axis piezo stage (Mad City Labs). Two 6 mm-diameter cylindrical shaped N52 magnets were affixed to a custom-fabricated aluminum bracket, which was attached to a 0.5”-travel XYZ translation stage equipped with a motorized Z-axis actuator (Thorlabs, Newton, New Jersey, USA). The stage position was homed before measurement by stepping the Z-actuator until the magnet bracket just touched the top of the sample. Measurements were performed while the Nikon Perfect Focus feedback system was turned on.

Samples were illuminated from the objective side via a mint green LED (Thorlabs) reflected into the back focal plane of a 100x Plan-Apo TIRF objective using a polarizing beamsplitter (Edmund Optics). Reflected cross-polarized light was transmitted through the beamsplitter and collected by an sCMOS camera (Hamamatsu Orca Flash v4). Bead images were collected using a custom-written LabView script or MicroManager 1.4 software^81^.

Magnetic tweezers force calibration was performed as described previously^82^ using transverse fluctuations of a magnetic bead tethered to a glass coverslip via lambda phage DNA (New England Biolabs).

#### Data analysis

A nonspecifically adhered bead was used to generate a reference image lookup table (LUT) while the piezo stage was stepped in increments of 10 nm. These LUTs were used to estimate the *z*-position of a query bead image. Briefly, the two-dimensional Fourier transform of the query bead image was calculated, and the radial profile extracted. The bead position was then estimated as follows: The one-dimensional Fourier transform of the LUT radial profiles was computed. The phase of the highest-amplitude frequency was then plotted against the bead height corresponding to the LUT slice, and a third-order polynomial was fit to this curve. This polynomial fit was then used to directly estimate height from the Fourier transform of the query image. Images were analyzed with both methods and found to produce similar results, reducing the chance that measured bead heights were analysis artifacts.

#### Sample preparation

SpyCatcher-ELP-functionalized #1.5 glass coverslips were prepared as previously described^83^, and stored in vacuum-sealed boxes at −20 °C. Briefly, 3-5 mm wide slits were cut with a razor blade into a piece of Parafilm M (Bemis) and sandwiched between a glass slide (1 mm thick, Fisher Scientific) and a SpyCatcher-ELP coverslip, to produce channels of ∼20 microliter volume. The assembly was heated on a 60 °C hot plate for 1 minute in increments of 15 seconds to adhere the parafilm to the glass. Channels were passivated overnight in blocking buffer (1 mg/mL Casein and 0.1% (v/v) Tween-20 in PBS). Samples were then washed with 10 channel volumes of assay buffer (50mM HEPES, 150 mM NaCl, pH 7.5) and incubated with 100 pM to 10 nM biotinylated fusion protein, for 1.5 hours. Samples were then washed with 10 channel volumes of buffer again, and then incubated with 1:10 M270-Streptavidin superparamagnetic beads diluted in blocking buffer for 25 minutes. Samples were then washed a final time with 10 channel volumes of assay buffer before measurement.

#### Force-jump experiments

Force jump experiments were performed as follows: A suitable bead tethered by a single complex was identified by comparing its force-extension behavior to that of the ELP, and centered in the microscope field of view. The magnetic tweezer was kept fully retracted from the coverslip, corresponding to sub-pN forces.

In each cycle, the magnet was quickly lowered to a set value while the bead height was monitored. Once the target height was reached, the magnet was held in place until the bead height exceeded a set threshold, at which point the magnet was then fully raised. Between cycles, the magnet was kept at maximum height for 30 seconds to allow the complex to return to binding equilibrium.

Extension traces over time for the force-jump cycles were analyzed as follows: a single-step profile with linear loading and unloading steps was fit with a least-squares objective function for each trace using the NumPy SHGO constrained global optimization routine, from which the bond lifetime and step size were measured. Traces were manually inspected to retain only those for which the pre-extension of the ELP handle was between 30 and 60 nm, and for which a step unbinding was clearly visible by eye.

#### Force-clamp experiments

Force-clamp experiments were performed following the force-jump protocol with the modification that instead of cycling the magnet position, the magnet position was held in a fixed position to obtain constant-force measurements.

#### Catch bond model fitting

Multiple kinetic models have been used to describe catch bonds^52^. Here, we noted that the PDZ-ligand dissociation kinetics measured for individual beads were, in general, well-described by single-exponential fits (Supp. Fig. S5, S6). Although this observation does not rule out more complex models^62,84^, relatively simpler models containing a single bound state, or else multiple bound states in rapid equilibrium, are thus adequate to describe the data. Of these, the two-pathway model performed better than the allosteric catch bond model^85^ as judged by the AIC. Both fits to the original data and bootstrap fitting to determine parameter errors were robust for both the nectin and JAM-A datasets. In addition, the rates *k_1_*, *k_2_*, and distance parameters *x_1_*, *x_2_*, were physically reasonable for both datasets. Neither the allosteric model^85^ nor the active-site deformation models^64^ met these criteria.

Bond lifetimes were fit using the genetic algorithm (`ga`) implemented in Matlab, such that the negative log likelihood function was minimized. The fits were constrained by requiring that the inferred dissociation rate at zero force match that measured using bio-layer interferometry. Multiple replicate fits were performed (1000 for the original data, 100 for each bootstrapped, synthetic dataset) to ensure convergence.

#### Steered Molecular Dynamics

We used the crystal structure of the PDZ domain of afadin in complex with the C-terminal peptide from nectin-3 (PDB 3AXA).^86^ To exhaustively sample the conformational landscape of the protein-peptide complex, the crystal structure was subjected to 1 microsecond of 4-replica implicit solvent MELD (Modeling Employed Limited Data) simulation^87,88^. A distribution of the non-bonded energy between afadin and nectin was computed, and representative structures from the most probable part of the distribution were selected for further thermalization in explicit solvent (Supp. Fig. S7B). Thereafter, the most tightly packed models were taken based on SASA distributions (Supp. Fig. S8B), which we take as the structure in the absence of force (Fig. 4A). These models were subjected to explicit solvent, biased SMD (Steered Molecular Dynamics) simulations^58,59^, where the C-terminal nitrogen atom of afadin was harmonically restrained and the C-terminal nitrogen atom of the five-residue nectin peptide was pulled away at a constant speed of 0.5 Å ns^-1^. The biased simulations were performed for 100 ns to enforce a C-terminal N atom separation by steps of 1.5 Å. Across 14 such steps, the protein-peptide distance changed from 4 – 20 Å, meaning that in total 1.4 microsecond simulations are performed. The “with force” conformation was extracted from the most probable region of the SASA distribution of the SMD run (Fig. 4B, Supp. Fig. S9).

