## Supplementary Material for "PDZ Domains from the Junctional Proteins Afadin and ZO-1 Act as Mechanosensors"

### Magnetic tweezer fusion construct sequences

*Afadin/JAM-A*

**Start** - 6x His linker Spytag linker Afadin PDZ long GS linker ybbR tag linker eGFP linker JAM-A ICD - **Stop**

**ATG**GGCAGCAGCCATCATCATCATCATCACAGCAGCGGCAGGGGCGGTGCCCACATCGTGATGGTGGACG
CCTACAAGCCCACCAAGGGGTCTAGCGGAGGTGCAGATTATGAAAGTCACCTTCTGCGTGAGAACACAGA
GCTGGCTCAGCCTCTGAGGAAAGAACCTGAAATAATCACTGTGACCCTAAAAAAGCAGAATGGAATGGGC
CTTAGCATTGTTGCAGCAAAGGGTGCTGGTCAAGATAAACTAGGAATCTACGTGAAGTCGGTTGTGAAAG
GAGGTGCTGCAGATGTGGATGGACGTCTAGCTGCAGGTGATCAGCTCCTCAGTGTGGATGGACGAAGTCT
GGTTGGACTCTCTCAGGAAAGGGCGGCAGAACTCATGACAAGAACAAGCTCTGTGGTGACACTGGAAGTA
GCAAAGCAGGGTGCCATCTACCACGGTCTGGCCACCCTTCTCAATCAGCCATCCCCCATGATGGGTGGTA
GCTCTGGTGAAAACCTGTATTTTCAGGGCGGTGGTTCTGCCGGTGGCTCCGGTTCTGGCTCCAGCGGTGG
CAGCTCTGGTGCGTCCGGCACGGGTACTGCGGGTGGCACTGGCAGCGGTTCCGGTACTGGCTCTGGCGGT
GGTTCTGGCGGCGGTTCTGAAGGTGGCGGCTCCGAAGGCGGCGGCAGCGAGGGCGGTGGTAGCGAAGGTG
GTGGCTCCGAGGGTGGCGGTTCCGGCGGCGGTAGCGGTGGATCGTCCGGAGAGAATCTTTATTTTCAGGG
CGAGCAGCGGCACTCTCTGGAATTCATCGCTTCTAAACTGGCTGGCAGCAGCATGGTGAGCAAGGGCGAG
GAGCTGTTCACCGGGGTGGTGCCCATCCTGGTCGAGCTGGACGGCGACGTAAACGGCCACAAGTTCAGCG
TGTCCGGCGAGGGCGAGGGCGATGCCACCTACGGCAAGCTGACCCTGAAGTTCATCTGCACCACCGGCAA
GCTGCCCGTGCCCTGGCCCACCCTCGTGACCACCCTGACCTACGGCGTGCAGTGCTTCAGCCGCTACCCC
GACCACATGAAGCAGCACGACTTCTTCAAGTCCGCCATGCCCGAAGGCTACGTCCAGGAGCGCACCATCT
TCTTCAAGGACGACGGCAACTACAAGACCCGCGCCGAGGTGAAGTTCGAGGGCGACACCCTGGTGAACCG
CATCGAGCTGAAGGGCATCGACTTCAAGGAGGACGGCAACATCCTGGGGCACAAGCTGGAGTACAACTAC
AACAGCCACAACGTCTATATCATGGCCGACAAGCAGAAGAACGGCATCAAGGTGAACTTCAAGATCCGCC
ACAACATCGAGGACGGCAGCGTGCAGCTCGCCGACCACTACCAGCAGAACACCCCCATCGGCGACGGCCC
CGTGCTGCTGCCCGACAACCACTACCTGAGCACCCAGTCCGCCCTGAGCAAAGACCCCAACGAGAAGCGC
GATCACATGGTCCTGCTGGAGTTCGTGACCGCCGCCGGGATCACTCTCGGCATGGACGAGCTGTACAAGG
GTAGCGGATCTGGTTCAGCGGCCGCATATAGACGGGGATACTTTGACCGGGCCAAGAAGGGCACCAGCAG
CAAGAAAGTGATCTACTCACAGCCCGCAGCACGAAGCGAAGGGGAGTTTAGGCAGACATCATCATTCCTG
GTG**TAA**

*Afadin/Nectin-1*

**Start** - 6x His linker Spytag linker Afadin PDZ long GS linker ybbR tag linker eGFP linker Nectin-1 ICD - **Stop**

**ATG**GGCAGCAGCCATCATCATCATCATCACAGCAGCGGCGGTAGGGGCGGTGCCCACATCGTGATGGTGG
ACGCCTACAAGCCCACCAAGGGGTCTAGCGGAGGTGCAGATTATGAAAGTCACCTTCTGCGTGAGAACAC
AGAGCTGGCTCAGCCTCTGAGGAAAGAACCTGAAATAATCACTGTGACCCTAAAAAAGCAGAATGGAATG
GGCCTTAGCATTGTTGCAGCAAAGGGTGCTGGTCAAGATAAACTAGGAATCTACGTGAAGTCGGTTGTGA
AAGGAGGTGCTGCAGATGTGGATGGACGTCTAGCTGCAGGTGATCAGCTCCTCAGTGTGGATGGACGAAG
TCTGGTTGGACTCTCTCAGGAAAGGGCGGCAGAACTCATGACAAGAACAAGCTCTGTGGTGACACTGGAA
GTAGCAAAGCAGGGTGCCATCTACCACGGTCTGGCCACCCTTCTCAATCAGCCATCCCCCATGATGGGTG
GTAGCTCTGGTGAAAACCTGTATTTTCAGGGCGGTGGTTCTGCCGGTGGCTCCGGTTCTGGCTCCAGCGG
TGGCAGCTCTGGTGCGTCCGGCACGGGTACTGCGGGTGGCACTGGCAGCGGTTCCGGTACTGGCTCTGGC
GGTGGTTCTGGCGGCGGTTCTGAAGGTGGCGGCTCCGAAGGCGGCGGCAGCGAGGGCGGTGGTAGCGAAG
GTGGTGGCTCCGAGGGTGGCGGTTCCGGCGGCGGTAGCGGTGGATCGTCCGGAGAGAATCTTTATTTTCA
GGGCGAGCAGCGGCACTCTCTGGAATTCATCGCTTCTAAACTGGCTGGCAGCAGCATGGTGAGCAAGGGC
GAGGAGCTGTTCACCGGGGTGGTGCCCATCCTGGTCGAGCTGGACGGCGACGTAAACGGCCACAAGTTCA
GCGTGTCCGGCGAGGGCGAGGGCGATGCCACCTACGGCAAGCTGACCCTGAAGTTCATCTGCACCACCGG
CAAGCTGCCCGTGCCCTGGCCCACCCTCGTGACCACCCTGACCTACGGCGTGCAGTGCTTCAGCCGCTAC
CCCGACCACATGAAGCAGCACGACTTCTTCAAGTCCTCCGCCATGCCCGAAGGCTACGTCCAGGAGCGCA
CCATCTTCTTCAAGGACGACGGCAACTACAAGACCCGCGCCGAGGTGAAGTTCGAGGGCGACACCCTGGT
GAACCGCATCGAGCTGAAGGGCATCGACTTCAAGGAGGACGGCAACATCCTGGGGCACAAGCTGGAGTAC
AACTACAACAGCCACAACGTCTATATCATGGCCGACAAGCAGAAGAACGGCATCAAGGTGAACTTCAAGA
TCCGCCACAACATCGAGGACGGCAGCGTGCAGCTCGCCGACCACTACCAGCAGAACACCCCCATCGGCGA
CGGCCCCGTGCTGCTGCCCGACAACCACTACCTGAGCACCCAGTCCGCCCTGAGCAAAGACCCCAACGAG
AAGCGCGATCACATGGTCCTGCTGGAGTTCGTGACCGCCGCCGGGATCACTCTCGGCATGGACGAGCTGT
ACAAGGGTAGCGGATCTGGTTCAGCGGCCGCAAGGAGACGGAGACATACTTTCAAGGGCGACTACTCTAC
CAAGAAGCACGTGTACGGCAACGGCTATAGCAAGGCAGGAATCCCACAGCACCACCCACCTATGGCCCAG
AATCTGCAGTACCCTGACGATAGCGACGATGAGAAGAAGCCAGGACCACTGGGAGGAAGCTCCTATGAGG
AAGAAGAAGAAGAAGAGGGAGGAGGAGGAGGCGAGAGGAAAGTGGGCGGCCCTCACCCAAAGTACGACGA
GGATGCCAAGCGGCCATACTTCACCGTGGATGAGGCAGAGGCCCGGCAGGACGGATATGGCGATAGAACA
CTGGGCTACCAGTATGACCCCGAGCAGCTGGATCTGGCCGAGAATATGGTGTCACAGAACGATGGGTCCT
TCATTAGCAAGAAAGAGTGGTACGTG**TAA**

*ZO-1/JAM-A*

**Start** - 6x His linker ZO-1 PSG linker Ybbr tag long GS linker eGFP linker Spytag linker JAM-A ICD - **Stop**

**ATG**GGCAGCAGCCATCATCATCATCATCACGGCAGCAGCATGCATATGATTCTTCGGCCCAGCATGAAAT
TGGTAAAATTCAGAAAAGGAGATAGTGTGGGTTTGCGGCTGGCTGGTGGAAATGATGTTGGAATATTTGT
AGCTGGCGTTCTAGAAGATAGCCCTGCAGCCAAGGAAGGCTTAGAGGAAGGTGATCAAATTCTCAGGGTA
AACAACGTAGATTTTACAAATATCATAAGAGAAGAAGCCGTCCTTTTCCTGCTTGACCTCCCTAAAGGAG
AAGAAGTGACCATATTGGCTCAGAAGAAGAAGGATGTTTATCGTCGCATTGTAGAATCAGATGTAGGAGA
TTCTTTCTATATTAGAACCCATTTTGAATATGAAAAGGAATCTCCCTATGGACTTAGTTTTAACAAAGGA
GAGGTGTTCCGTGTTGTGGATACCTTGTACAATGGAAAACTGGGCTCTTGGCTTGCTATTCGAATTGGTA
AAAATCATAAGGAGGTAGAACGAGGCATCATCCCTAATAAGAACAGAGCTGAGCAGCTAGCCAGTGTACA
GTATACACTTCCAAAAACAGCAGGCGGAGACCGTGCTGACTTCTGGAGATTCAGAGGTCTTCGCAGCTCC
AAGAGAAATCTTCGAAAAAGCAGAGAGGATTTGTCCGCTCAGCCTGTTCAAACAAAGTTTCCAGCTTATG
AAAGAGTGGTTCTTCGAGAAGCTGGATTTCTGAGGCCTGTAACCATTTTTGGACCAATAGCTGATGTTGC
CAGAGAAAAGCTGGCAAGAGAAGAACCAGATATTTATCAAATTGCAAAGAGTGAACCACGAGACGCTGGA
ACTGACCAACGTAGCTCTGGCATTATTCGCCTGCATACAATAAAGCAAATCATAGATCAAGACAAGCATG
CATTATTAGATGTAACACCAAATGCAGTTGATCGTCTTAACTATGCCCAGTGGTATCCAATTGTTGTATT
TCTTAACCCTGATTCTAAGCAAGGAGTAAAAACAATGAGAATGAGGTTATGTCCAGAATCTCGGAAAAGT
GCCAGGAAGTTATACGAGCGATCTCATAAACTTCGTAAAAATAATCACCATCTTTTTACAACTACAATTA
ACTTAAATTCAATGAATGATGGTTGGTATGGTGCGCTGAAAGAAGCAATTCAACAACAGCAAAACCAGCT
GGTATGGGTTTCCGAGGGAAGCAGCGGCGGTAGGGGCGGTGACTCTCTGGAATTCATCGCTTCTAAACTG
GCTGGTGGTAGCTCTGGTGAAAACCTGTATTTTCAGGGCGGTGGTTCTGCCGGTGGCTCCGGTTCTGGCT
CCAGCGGTGGCAGCTCTGGTGCGTCCGGCACGGGTACTGCGGGTGGCACTGGCAGCGGTTCCGGTACTGG
CTCTGGCGGTGGTTCTGGCGGCGGTTCTGAAGGTGGCGGCTCCGAAGGCGGCGGCAGCGAGGGCGGTGGT
AGCGAAGGTGGTGGCTCCGAGGGTGGCGGTTCCGGCGGCGGTAGCGGTGGATCGTCCGGAGAGAATCTTT
ATTTTCAGGGCGAGCAGCGGGTGAGCAAGGGCGAGGAGCTGTTCACCGGGGTGGTGCCCATCCTGGTCGA
GCTGGACGGCGACGTAAACGGCCACAAGTTCAGCGTGTCCGGCGAGGGCGAGGGCGATGCCACCTACGGC
AAGCTGACCCTGAAGTTCATCTGCACCACCGGCAAGCTGCCCGTGCCCTGGCCCACCCTCGTGACCACCC
TGACCTACGGCGTGCAGTGCTTCAGCCGCTACCCCGACCACATGAAGCAGCACGACTTCTTCAAGTCCGC
CATGCCCGAAGGCTACGTCCAGGAGCGCACCATCTTCTTCAAGGACGACGGCAACTACAAGACCCGCGCC
GAGGTGAAGTTCGAGGGCGACACCCTGGTGAACCGCATCGAGCTGAAGGGCATCGACTTCAAGGAGGACG
GCAACATCCTGGGGCACAAGCTGGAGTACAACTACAACAGCCACAACGTCTATATCATGGCCGACAAGCA
GAAGAACGGCATCAAGGTGAACTTCAAGATCCGCCACAACATCGAGGACGGCAGCGTGCAGCTCGCCGAC
CACTACCAGCAGAACACCCCCATCGGCGACGGCCCCGTGCTGCTGCCCGACAACCACTACCTGAGCACCC
AGTCCGCCCTGAGCAAAGACCCCAACGAGAAGCGCGATCACATGGTCCTGCTGGAGTTCGTGACCGCCGC
CGGGATCACTCTCGGCATGGACGAGCTGTACAAGGGTAGCGGATCTGGTTCAGCCCACATCGTGATGGTG
GACGCCTACAAGCCCACCAAGGGGTCTAGCGGAGCGGCCGCATATAGACGGGGATACTTTGACCGGGCCA
AGAAGGGCACCAGCAGCAAGAAAGTGATCTACTCACAGCCCGCAGCACGAAGCGAAGGGGAGTTTAGGCA
GACATCATCATTCCTGGTG**TAA**

### Supplementary Figures

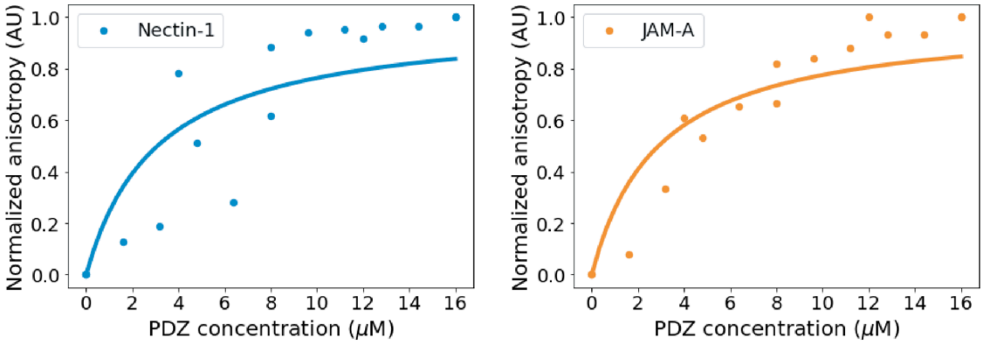


**Figure S1.** Fluorescence anisotropy measurements of PDZ-peptide binding between the soluble Afadin PDZ domain and FITC-labeled Nectin-1 intracellular peptide (*left*) and JAM-A intracellular peptide (*right*). Normalized fluorescence anisotropies for *N* = 2 experimental days are plotted. Lines indicate best-fit Langmuir binding isotherms.

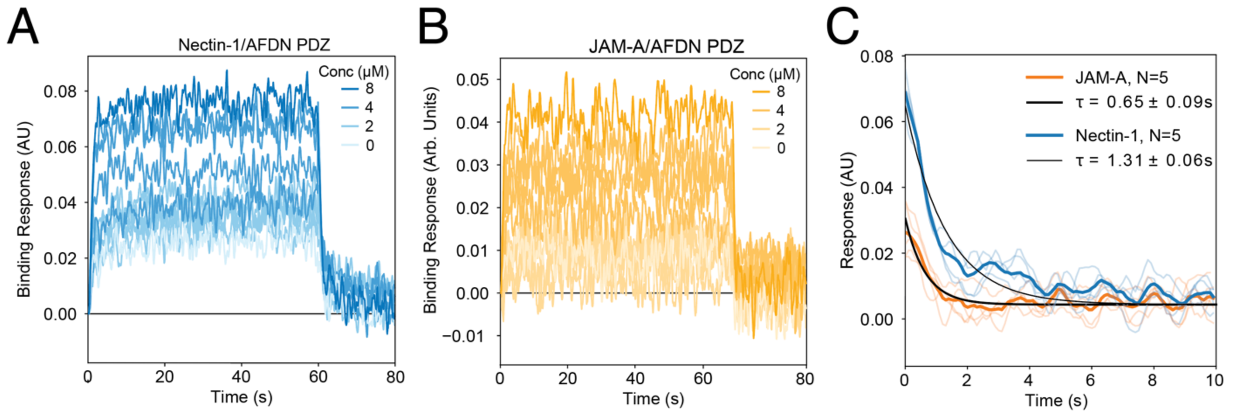


**Figure S2.** (A) All BLI traces for nectin-1/afadin PDZ BLI measurements from Figure 2A. (B) All BLI traces for JAM-A/afadin PDZ from Figure 2B. (C) BLI traces and curve fits of 4 μM Afadin dissociation from the ICD of nectin-1 (*blue*) and JAM-A (*orange*).

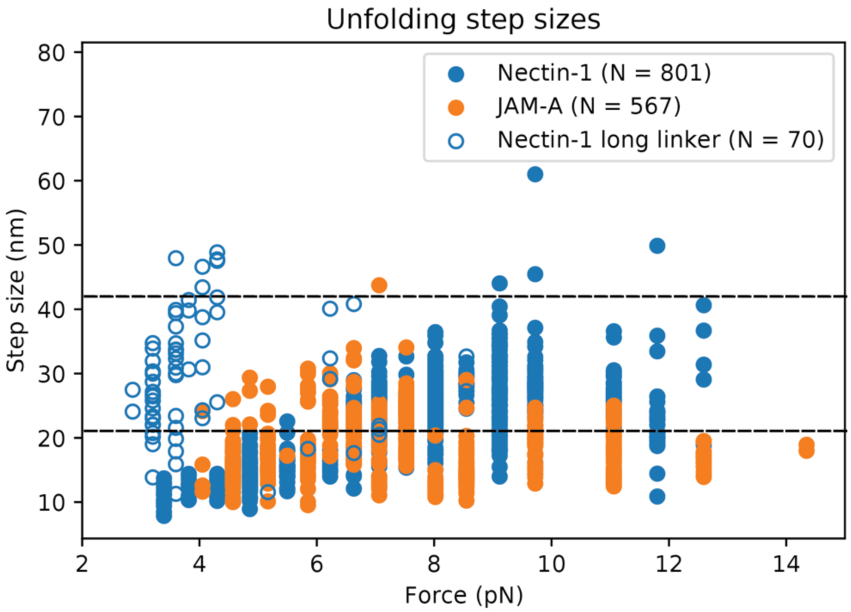


**Figure S3.** Step sizes measured in afadin-PDZ force-jump experiments as a function of applied force. Open circles denote single-length linker constructs, open circles denote double-length constructs. Expected contour length for the double-length linker is 42 nm (dashed line).

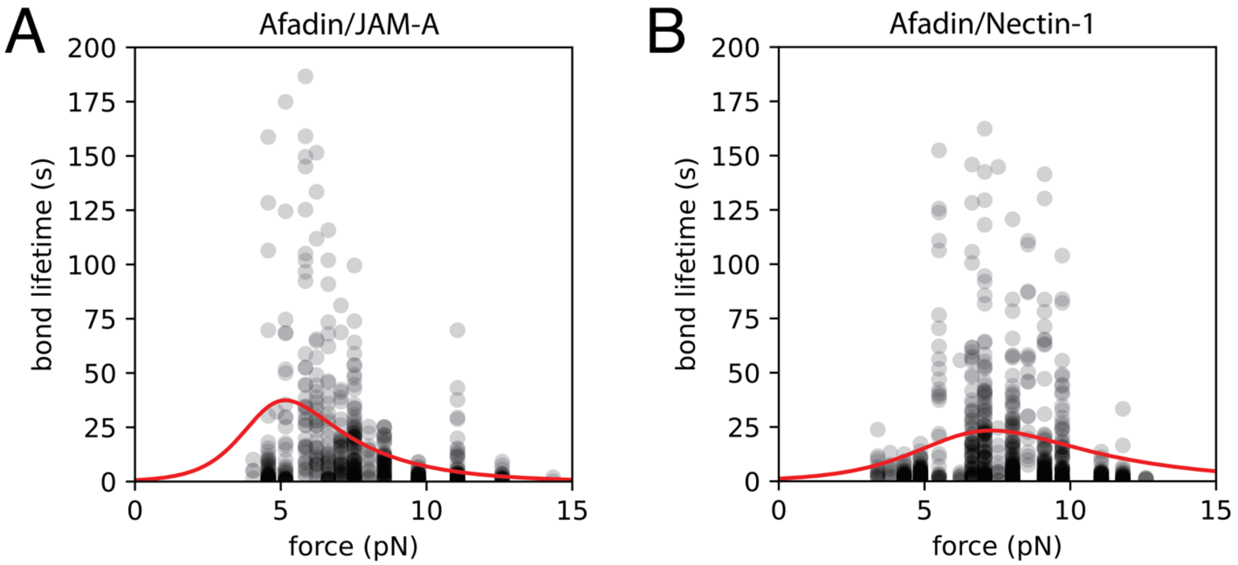


**Fig. S4**. Two-pathway catch bond fits overlaid with individual data points measured for the afadin/JAM-A (A) and afadin/nectin-1 (B) constructs.

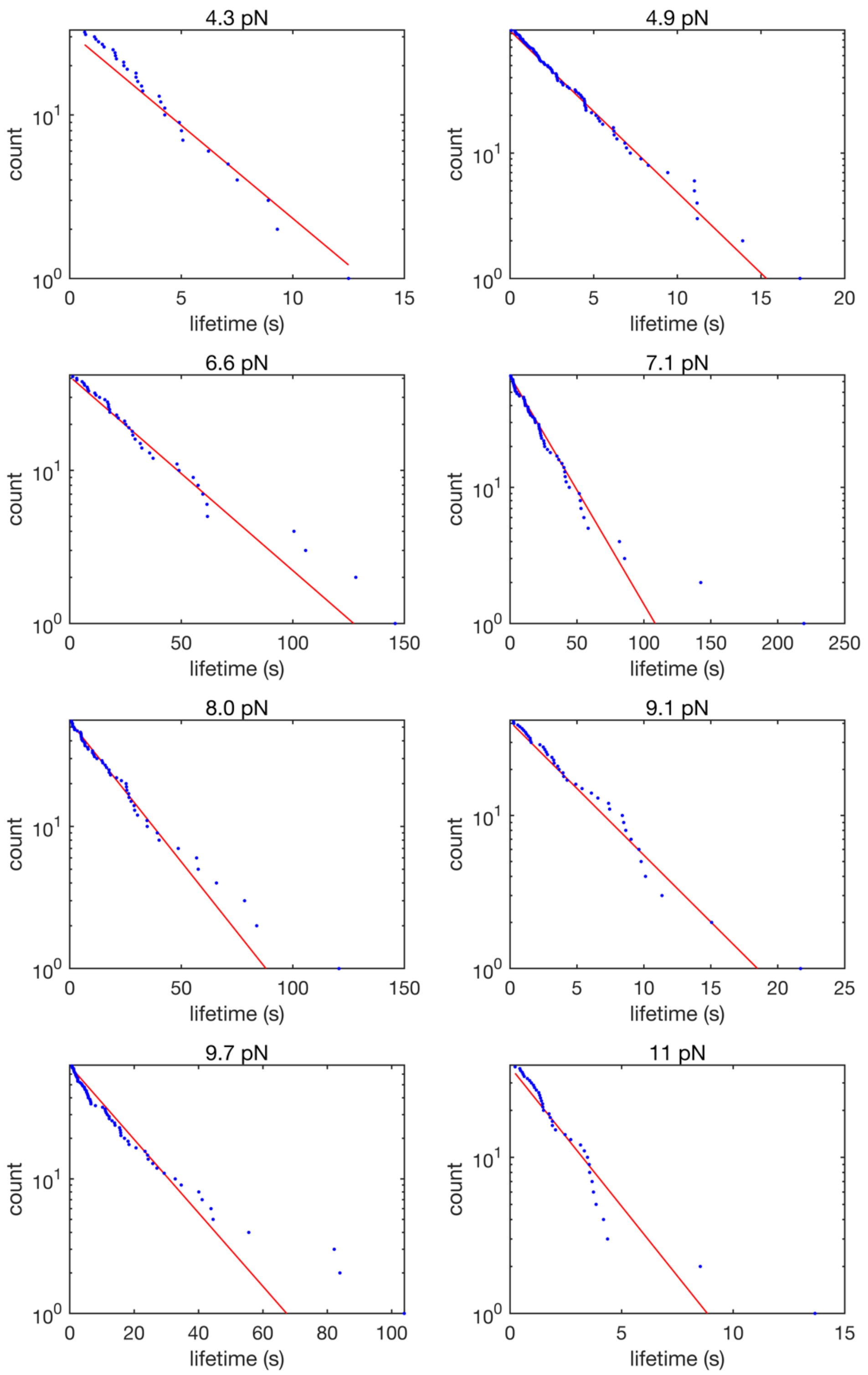


**Fig. S5**. Complementary cumulative distributions for afadin/nectin-1 bond lifetimes as a function for force. Each plot represents data collected from a single bead, with N > 30 observations for each bead.

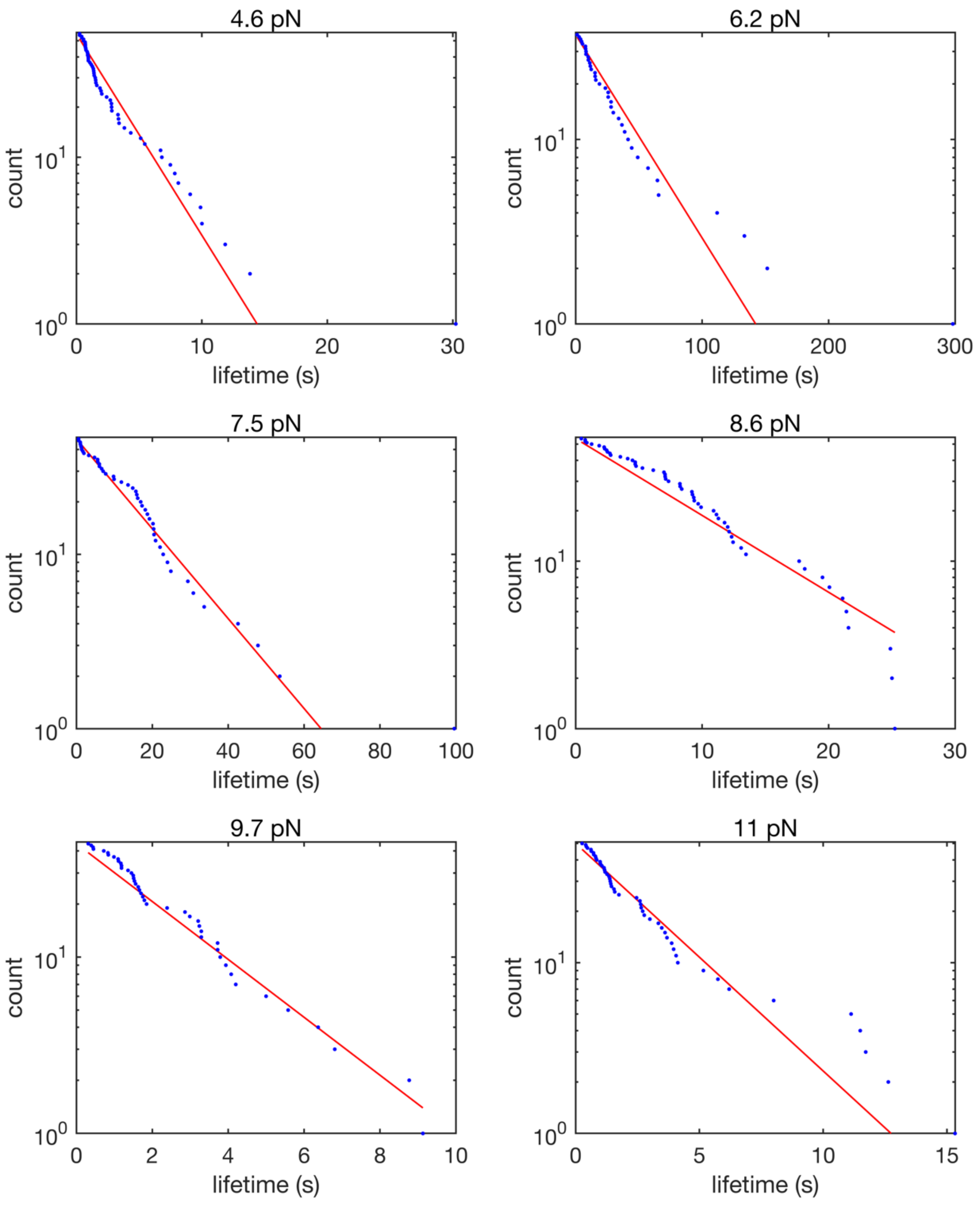


**Fig. S6**. Complementary cumulative distributions for afadin/JAM-A bond lifetimes as a function for force. Each plot represents data collected from a single bead, with N > 25 observations for each bead.

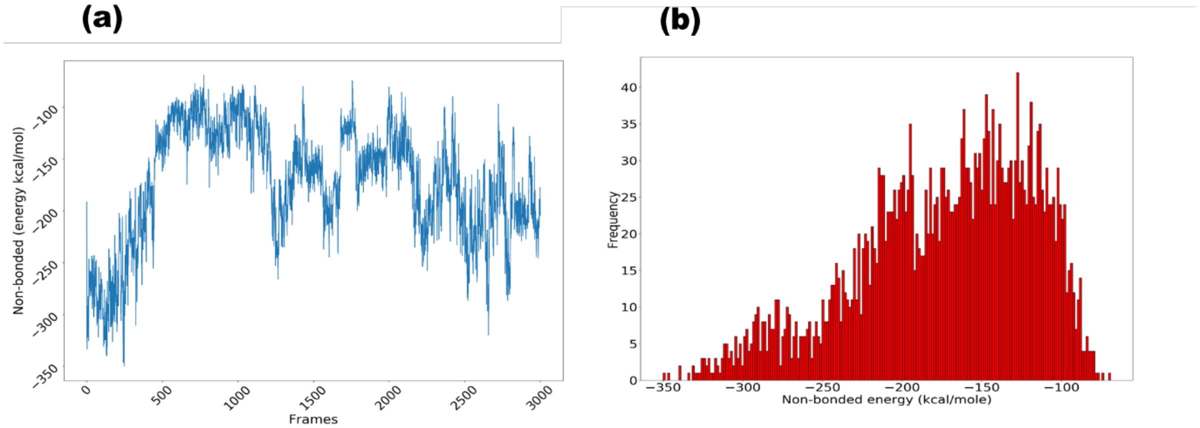


**Fig. S7**. (A) Non-bonded energy between nectin and afadin from the initial MELD simulation. (B) Non-bonded energy histogram. A representative structure was chosen from the most probable region for explicit solvent equilibration.

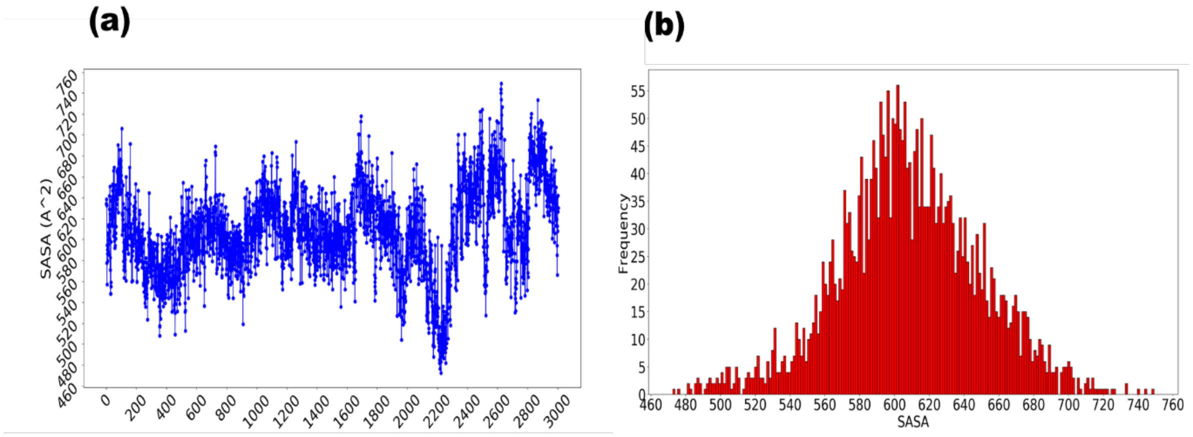


**Fig. S8**. (A) Solvent Accessible Surface Area (SASA) between nectin and afadin from the initial explicit solvent equilibration. (B) SASA histogram. The representative “without force” conformation was chosen from the most probable region.

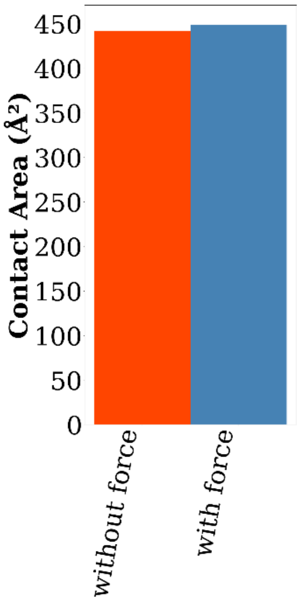


**Fig. S9**. Contact area plot between structures “with force” and “without force.” Contact area was calculated according to the formula:

Contact area = {(SASA of only nectin chain) + (SASA of the binding site of the PDZ domain of afadin)} – {SASA of the nectin chain and the PDZ domain of afadin}
